# Gene Expression and Splicing QTL Analysis of Blood Cells in African American Participants from the Jackson Heart Study

**DOI:** 10.1101/2023.04.26.538455

**Authors:** Jia Wen, Quan Sun, Le Huang, Lingbo Zhou, Margaret F. Doyle, Lynette Ekunwe, Nels C. Olson, Alexander P. Reiner, Yun Li, Laura M. Raffield

## Abstract

Most gene expression and alternative splicing quantitative trait loci (eQTL/sQTL) studies have been biased toward European ancestry individuals. Here, we performed eQTL and sQTL analysis using TOPMed whole genome sequencing-derived genotype data and RNA sequencing data from stored peripheral blood mononuclear cells in 1,012 African American participants from the Jackson Heart Study (JHS). At a false discovery rate (FDR) of 5%, we identified 4,798,604 significant eQTL-gene pairs, covering 16,538 unique genes; and 5,921,368 sQTL-gene-cluster pairs, covering 9,605 unique genes. About 31% of detected eQTL and sQTL variants with a minor allele frequency (MAF) > 1% in JHS were rare (MAF < 0.1%), and therefore unlikely to be detected, in European ancestry individuals. We also generated 17,630 eQTL credible sets and 24,525 sQTL credible sets for genes (gene-clusters) with lead QTL p < 5e-8. Finally, we created an open database, which is freely available online, allowing fast query and bulk download of our QTL results.

## Main

Quantitative trait loci (QTL) analysis of molecular phenotypes, such as gene expression levels (eQTLs) or alternative splicing events (sQTLs), can contribute to understanding the regulatory impact of individual genetic variants and the regulation of gene expression across the genome in a given tissue. Cataloging eQTL and sQTL variants across tissues can also help to functionally characterize non-coding genetic variants which constitute the majority of variants associated with complex diseases and traits through GWAS ^1–3^. Therefore, eQTL and sQTL analyses have been widely conducted to link genetic variants associated with complex traits and diseases to their likely target genes and underlying mechanisms ^3,4^.

Most eQTL/sQTL studies focus on European ancestry (EUR) individuals and few include substantial representation of African ancestry (AFR) individuals (based on similarity to reference panels such as 1000G). For example, the most commonly used eQTL/sQTL resources in blood are from GTEx (n=670, ~12% AFR individuals in v8) ^3^ and, for eQTL only, the eQTLGen meta-analysis including 31,684 participants (none AFR) ^5^. There have been a few recent efforts to include a greater number of AFR participants ^1,6–9^, but more efforts are still needed to match up with the sample size available in QTL studies for EUR individuals. Some of these existing studies in AFR individuals either utilize transformed cell lines (pre-publication African Functional Genomics Resource (AFGR) resource) rather than native tissue, rely on older microarray data ^7,8^ to measure gene expression (and therefore do not include examination of splicing QTLs), or genotyping array and imputation (rather than WGS) ^7,8^ for genotyping, or include only children ^9^ (and therefore may be less relevant for identifying mechanisms for genome-wide association study (GWAS) identified loci for complex traits or chronic diseases that mainly occur among adults).

To address this gap, we performed eQTL and sQTL analysis in 1,012 AFR individuals from the Jackson Heart Study (JHS) ^10^ with RNA-seq measured gene expression levels from stored peripheral blood mononuclear cells (PBMCs) and genome-wide genotyping from whole genome sequencing (WGS) generated through the NHLBI Trans-Omics for Precision Medicine (TOPMed) program. JHS participants are part of a longitudinal population-based cohort study focusing on improved understanding of risk factors for cardiometabolic disease among African Americans, including genetic risk factors. All participants have significant portions of their genomes with genetic similarity to African (AFR) reference populations, with mean and median African global ancestry component 83.4% and 85.4% (**Supplemental Materials**). To our knowledge, this is the largest sequencing-based eQTL/sQTL study to date in predominantly AFR populations. More details on cohort and methods are described in the **Supplemental Materials**.

As displayed in **Figure S1**, RNA-seq data were generated at the University of Washington Northwest Genomics Center (NWGC) at an average read depth of 50M utilizing stored PBMC obtained from 1,027 related JHS participants recruited from families residing in Jackson, Mississippi. We performed sample-level quality control (QC) by removing low RNA quality samples, and those with genotype inconsistencies between RNA and DNA based on verifyBamID ^11^ results, leaving 1,012 individuals (n=706 unrelated individuals with kinship < 0.2 from PC-AiR ^12^ pairwise kinship coefficient) for downstream analysis. Variant- and gene-level QC was then performed based on these 1,012 individuals by removing variants with minor allele frequency (MAF) < 1% and low expression genes (**Supplemental Materials**), resulting in 15,474,937 variants and 17,383 genes for eQTL analysis. For sQTL analysis, we adopted LeafCutter to calculate intron excision ratios to quantify alternative splicing events ^13^ and applied a QC strategy similar to GTEx ^3^ (**Supplemental Materials**), retaining 97,840 intron clusters for our sQTL analysis. We performed *cis*-eQTL and *cis-*sQTL analysis using linear mixed models implemented in APEX ^14^. The genetic relationship matrix (GRM), included as a random effect in the linear mixed model to account for relatedness, was calculated based on common variants (MAF > 1%) using GCTA ^15,16^. In the eQTL analysis, we inverse-normalized gene expression values and then calculated probabilistic estimations of expression residuals (PEER) factors ^17^. Covariates included as fixed effects in the linear models were age, sex, top 10 genotype PCs calculated in JHS samples using PC-AiR ^12^, and 70 PEER factors. The number of PEER factors was chosen based on the point where we no longer observed a substantive increase in the number of detected QTLs with the addition of an additional 10 PEER factors, for eQTL analysis (**Figure S4A**). For splicing QTL analyses, we adjusted for 25 splicing PCs, which maximized the significant variant-cluster pairs (**Figure S4B**).

Our eQTL analysis revealed 4,798,604 variant-gene pairs at an MAF >1% representing 16,538 unique genes at a genome-wide Benjamini-Hochberg-corrected FDR of 5%. We compared our eQTL results with those from 3 other blood or LCL-based published sources that include AFR ancestry individuals, GTEx v8 ^3^ (12.3% African American, total n=670), GENOA ^7^ (AA, n=1,032 related), and the AFGR study (n=593) (**Figure 1, S2, S3, Table S1**). Our results revealed a greater number of significant variant-gene pairs at the same MAF threshold and 17-50% increase in the number of genes with eQTLs (e-genes) compared to the other studies. The greater statistical power for QTL detection is likely due to the comparable or larger effective sample size and the genome and RNA sequencing versus array-based data in JHS. We then compared eQTL-gene association results by examining agreement in terms of either estimated effect size (beta) or statistical significance (p-value) (**Figure 1, S2, S3**). For every variant-gene pair shared between JHS and the previously published AFR blood eQTL datasets, we observed reasonable agreement with the correlation of effect sizes ranging from 0.49-0.82 (**Figure 1A, S2A, S3A**). By comparing p-values, we found that our JHS eQTLs were generally more significant compared to the other studies (**Figure 1B, S2B, S3B**). We also compared the agreement with restriction to the index variants for each gene, and observed similar consistencies in effect sizes (**Figure 1C, S2C, S3C**) and greater statistical significance (**Figure 1D, S2D, S3D**).

**Figure 1.**
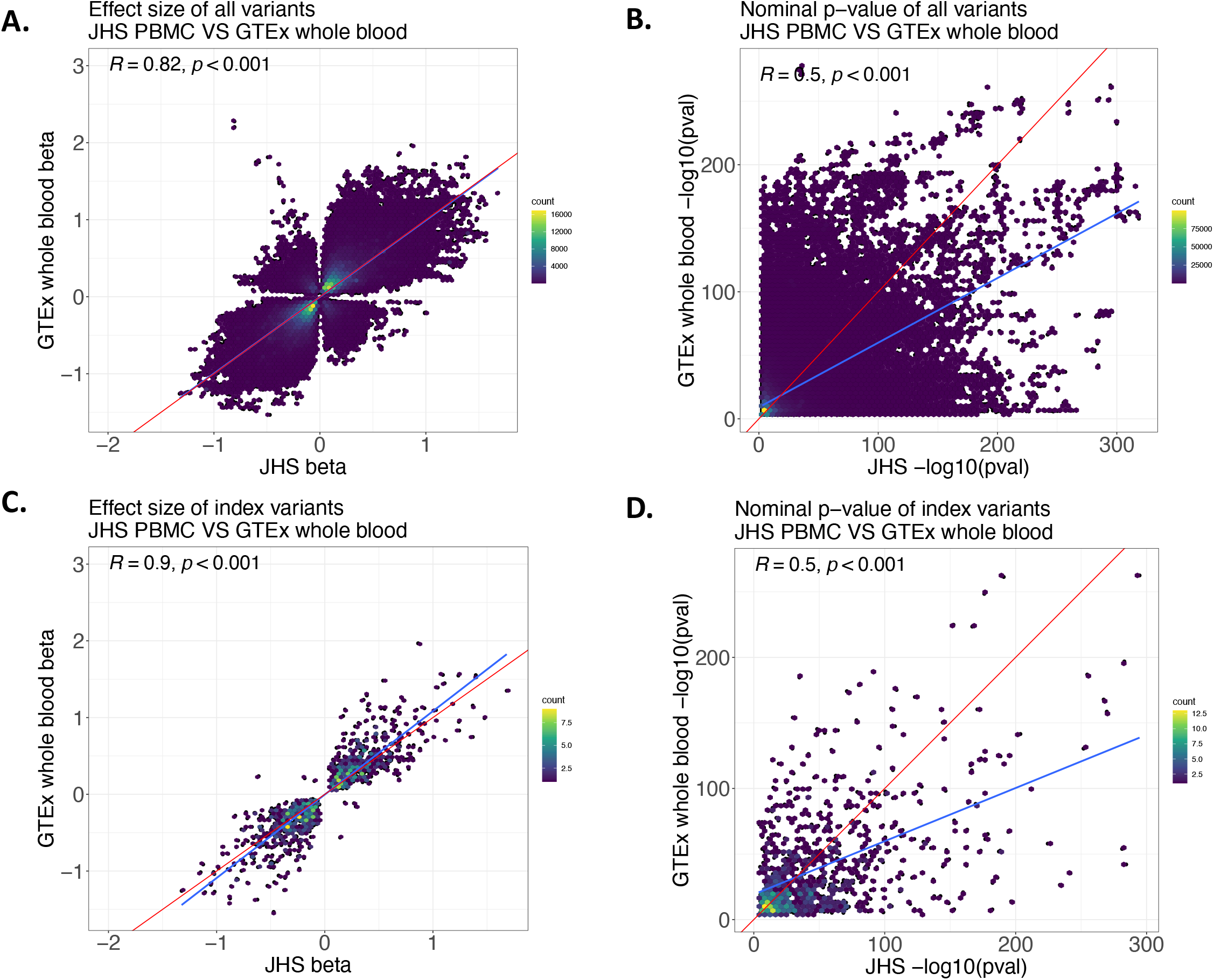
eQTL results comparison with GTEx whole blood eQTLs. **A and B.** Comparison of effect size estimates (**A**) and nominal p-values (**B**), for all shared variants. **C and D.** Comparison of effect size estimates (**C**) and nominal p-values (**D**) restricting to the index variants (i.e., the most significant eQTL variant for each gene). The red line denotes the diagonal line and the blue line is the regression fitting line.

Our sQTL results revealed 5,918,395 variant-gene-cluster pairs at FDR 5%, involving 9,605 unique genes (**Figure 2, Table S2**). There are fewer sQTL studies including AFR participants than eQTL studies, emphasizing the value of our study in JHS. In terms of unique genes with sQTLs, we achieved a 2.2-fold increase compared to GTEx v8. Since no gene information (only intron clusters) was provided in the AFGR pre-publication sQTL results, we could not compare to AFGR at gene level. We also compared the effect size estimates and p-values between our sQTL results and GTEx v8. **Figure 2** shows that the effect size estimates were in high agreement across all tested scenarios including shared variant-cluster pairs, index-variant-cluster pairs, and index-variant-gene pairs (R=0.79-0.89). In terms of statistical significance (based on p-values), our sQTLs were in general more significant than GTEx v8 sQTLs, again likely due to the increased sample size of JHS compared to GTEx.

**Figure 2.**
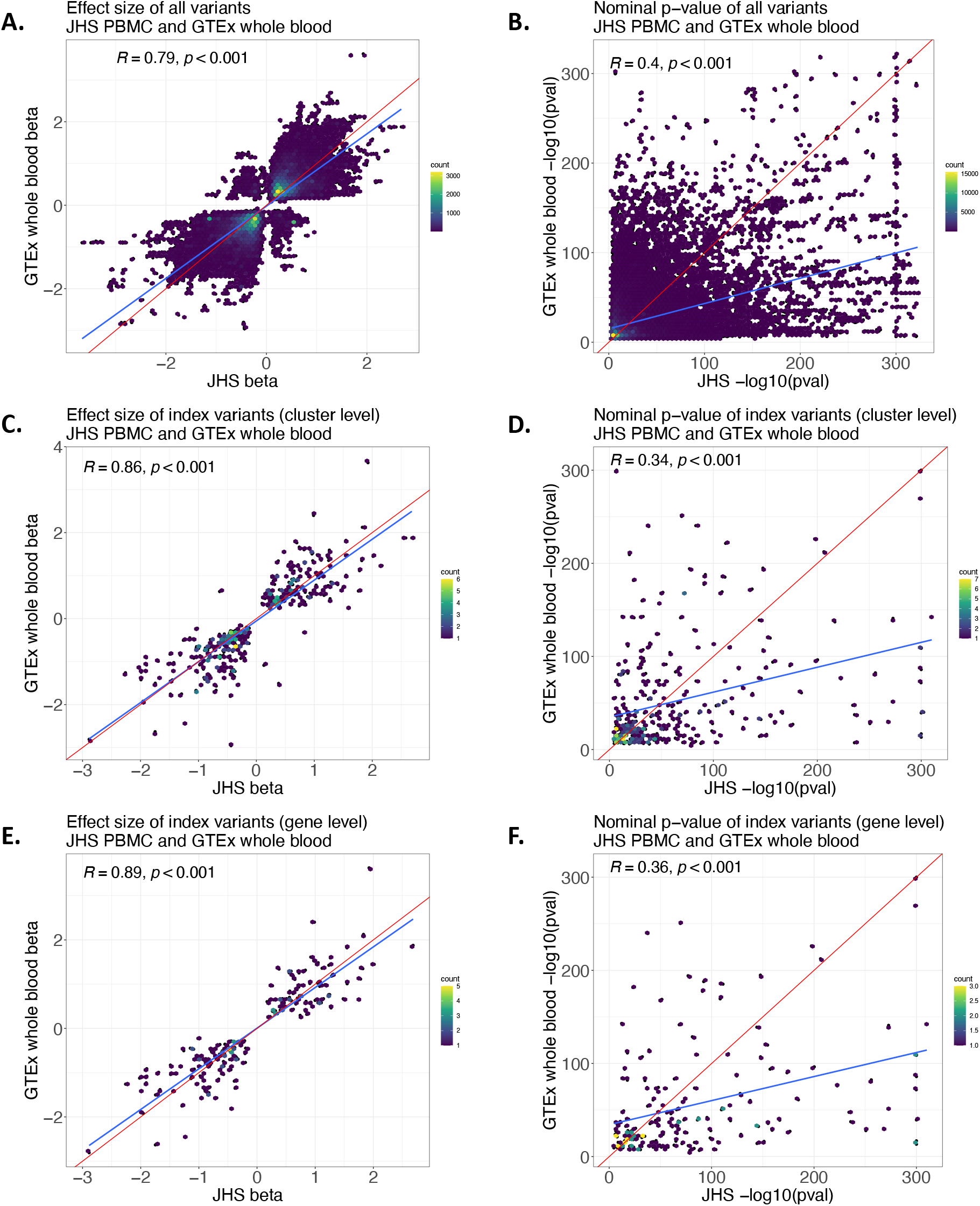
sQTL results comparison with GTEx whole blood sQTLs. **A and B.** Comparison of effect size estimates (**A**) and nominal p-values (**B**), for all shared variants. **C and D.** Comparison of effect size estimates (**C**) and nominal p-values (**D**) restricting to the cluster-level index variants (i.e., the most significant sQTL variant for each gene-cluster). **E and F.** Comparison of effect size estimates (**E**) and nominal p-values (**F**) restricting to the gene-level index variants (i.e., the most significant sQTL variant for each gene among all its clusters). The red line denotes the diagonal line and the blue line is the regression fitting line.

We next performed fine-mapping using SuSie ^18^ to identify and provide distinct eQTL and sQTL signals and the likely set of variants driving each signal. Fine mapping results can be valuable for GWAS integration and colocalization analyses with clinical phenotypes ^19,20^.

Out of the 9,678 genes with lead eQTL p-value < 5e-8, we generated credible sets for 9,317 genes with the remaining genes having an empty credible set, likely due to SuSie non-convergence. The vast majority (90.7%) of genes have <=50 total variants in their credible sets (**Figure 3A**) and most genes have a small number of distinct credible sets (**Figure 3C**).

**Figure 3.**
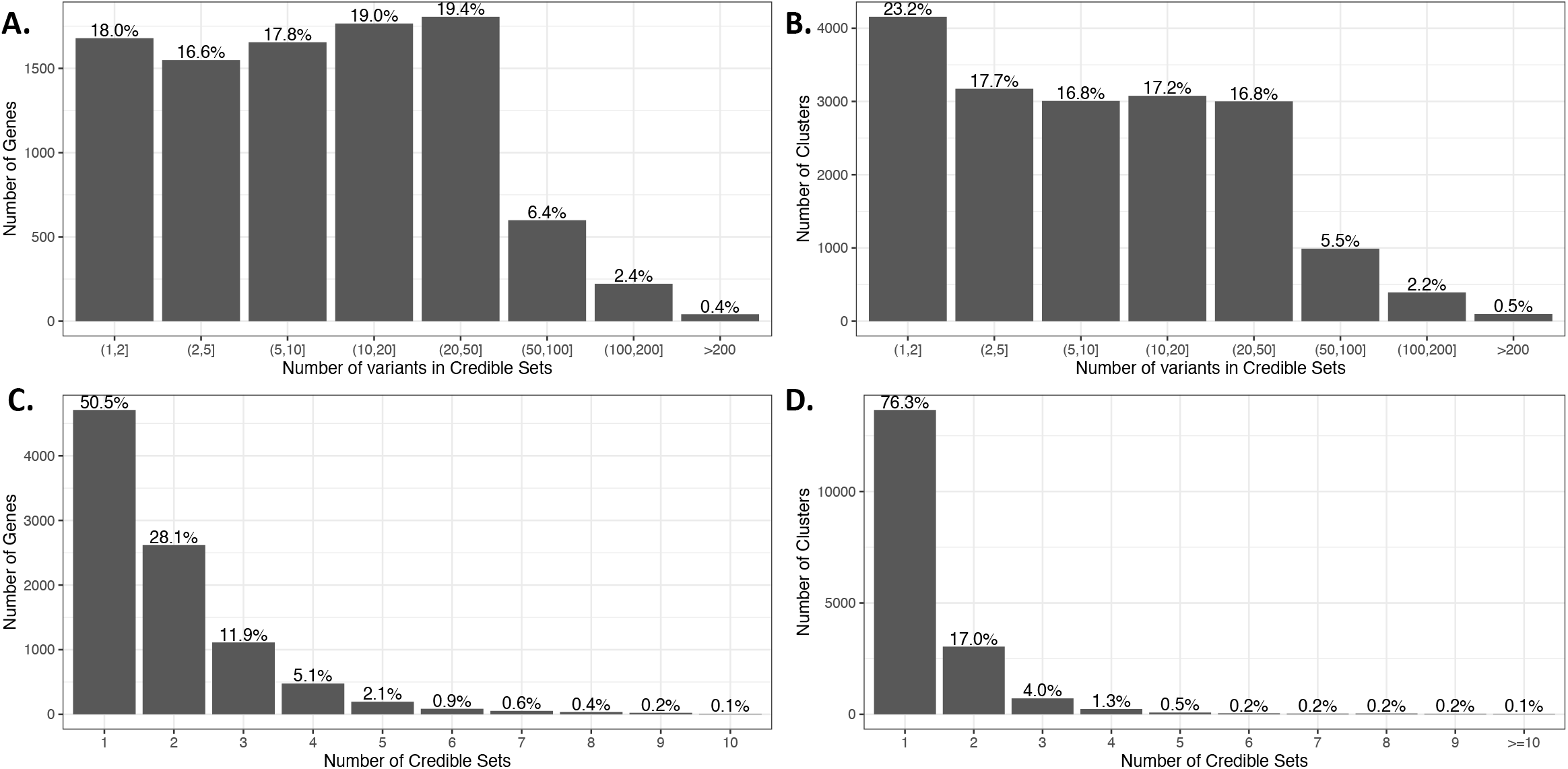
Histograms of SuSie fine-mapping results. **A**. Number of variants in credible sets for eQTL fine-mapping. **B.** Number of variants in credible sets for sQTL fine-mapping. **C**. Number of credible sets for eQTL fine-mapping. **D.** Number of credible sets for sQTL fine-mapping.

For example, 8,439 (90.6%) genes have <=3 credible sets while only 68 (0.7%) genes have >=8 credible sets. Similarly, most genes (94.5%) have 3 or less distinct signals in GTEx conditional analyses. Comparing the total number of distinct signals with GTEx, for genes with both a GTEx and a JHS significant signal, our results generated ~16K additional credible sets that are not present in GTEx (**Figure S5**). Conversely among the genes where SuSie fine-mapping was performed in JHS, there were about 10K unique GTEx distinct signals that were not identified in JHS. Similarly, we performed fine-mapping analyses for sQTLs at gene-cluster level for those clusters with lead sQTL p-value < 5e-8, resulting in 18,785 clusters representing 5,124 unique genes. We successfully generated credible sets for 17,896 clusters representing 5,038 unique genes, and the results show similar patterns as eQTLs in terms of number of distinct credible sets and number of variants within each set (**Figure 3B and 3D**).

Our eQTL and sQTL fine-mapping results can be utilized in analyses to link GWAS variants to genes ^21^. For example, in a multi-ancestry GWAS ^22^, rs10173412, an intronic variant of *RBMS1*, was a lead variant associated with lymphocyte counts (**Figure 4A**). rs10173412 is also in the primary credible set of *RBMS1* eQTLs (**Figure 4B**), suggesting potential regulatory links between variant to gene. Another example is rs113315762 (an intergenic variant located between *HRK* and *FBXW8)* associated with eosinophil counts (**Figure 4C**). In JHS, this variant is not the lead eQTL (**Figure 4D**); rather, rs113315762 is in the secondary credible set of *FBXW8* from our fine-mapping results. We further note that *HRK* has credible sets where this variant rs113315762 is not present, suggesting a potential regulatory role of the eosinophil-associated variant on blood cell *FBXW8* expression. In both of these examples, the two index trait-associated variants, rs10173412 and rs113315762, are more frequent in African ancestry populations than European ancestry (EUR v.s. AFR MAF 20.8% v.s. 32.4%, and 10.7% v.s. 45.9%, respectively ^23^). Moreover, the 12q24 region harboring *FBXW8* was recently identified as an African ancestry-specific risk loci for eosinophilic esophagitis in a genome-wide admixture and association analysis in African Americans ^24^. Neither the rs10173412-*RBMS1* nor rs113315762-*FBXW8* eQTLs are reported in GTEx v8 in any tissue, or tagged by any LD proxy. Therefore, these examples demonstrate the value of our results for interpreting and prioritizing GWAS variants more common in AFR and, as in prior eQTL/GWAS integration analyses ^20^, shows that additional insights can be gained through consideration of non-primary eQTL/sQTL signals.

**Figure 4.**
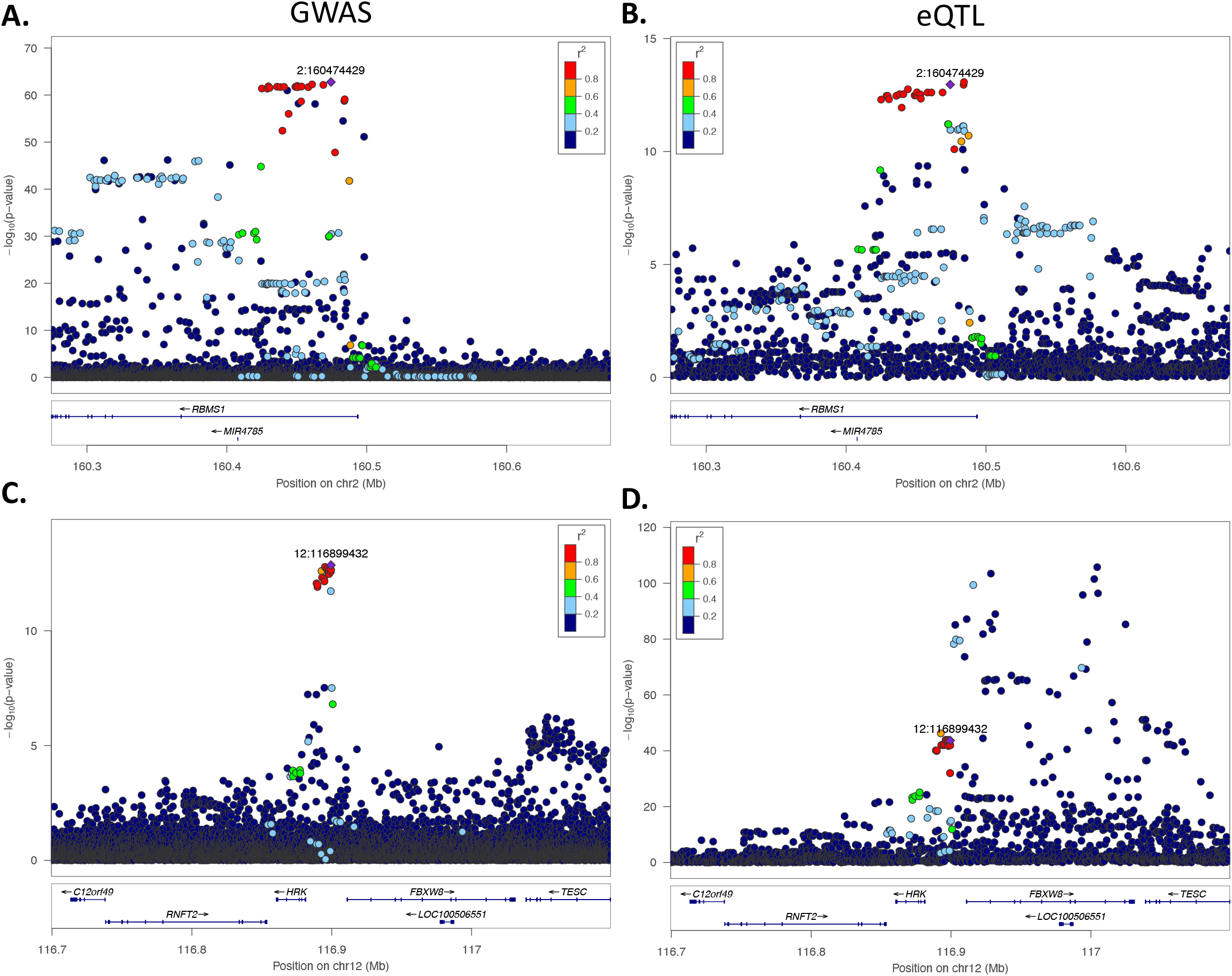
Locuszoom plot for overlapping GWAS and eQTL signals for two hematological trait GWAS signals. **A and C** show the GWAS associations; **B and D** show the eQTL associations**. A and B.** The example of rs10173412-*RBMS1*. **A.** Association of rs10173412 (2:160474429, hg38) with lymphocyte counts in a multi-ancestry GWAS study ^22^; **B.** association of rs10173412 (2:160474429, hg38) with *RBMS1* expression. **C and D.** The example of rs113315762-*FBXW8*. **C**. Association between rs113315762 (12:116899423) and eosinophil counts in the same GWAS study; **D**. association between rs113315762 (12:116899423) and expression of *FBXW8*. The genome build is hg38. For both GWAS and eQTL associations, LD is based on TOP-LD ^23^ AFR.

Although we demonstrate general agreement in effect directions and magnitude of associations for variant-gene and index-variant-gene pairs shared between our JHS analyses and previously published eQTL/sQTL datasets, some differences still exist. These differences may be due to several factors, such as MAF differences between AFR and EUR, demographic and environmental differences between populations, or technical differences in sample collection, storage, and library preparation, as well as statistical chance, especially in the modest sample sizes currently available for eQTL/sQTL analysis. Indeed, there are examples of QTL with large MAF differences between EUR and AFR, which would cause some AFR ancestry-specific QTLs to be missed in prior euro-centric studies. Overall, in our eQTL and sQTL results at FDR < 5%, 31.0% of eQTL variants and 30.7% of sQTL variants were rare (MAF < 0.1%) and 37.2% and 36.8% were low frequency (MAF < 0.5%) in EUR individuals, while all of these variants have MAF > 1% in JHS (**Figure 5**). These variants represent 4,272 eQTL and 5,830 sQTL credible sets with MAF EUR < 0.1%, and represent 5,240 eQTL and 7,124 sQTL credible sets with MAF EUR < 0.5%. Besides genetic ancestry difference in study subjects, GTEx utilized whole blood including cell types such as erythrocytes, platelets, and neutrophils while our JHS RNA-seq data was based on cryopreserved PBMCs which mainly consists of lymphocytes (T cells, B cells, and NK cells) and monocytes. Thus, differential gene expression by cell types can also potentially contribute to discrepant results.

**Figure 5.**
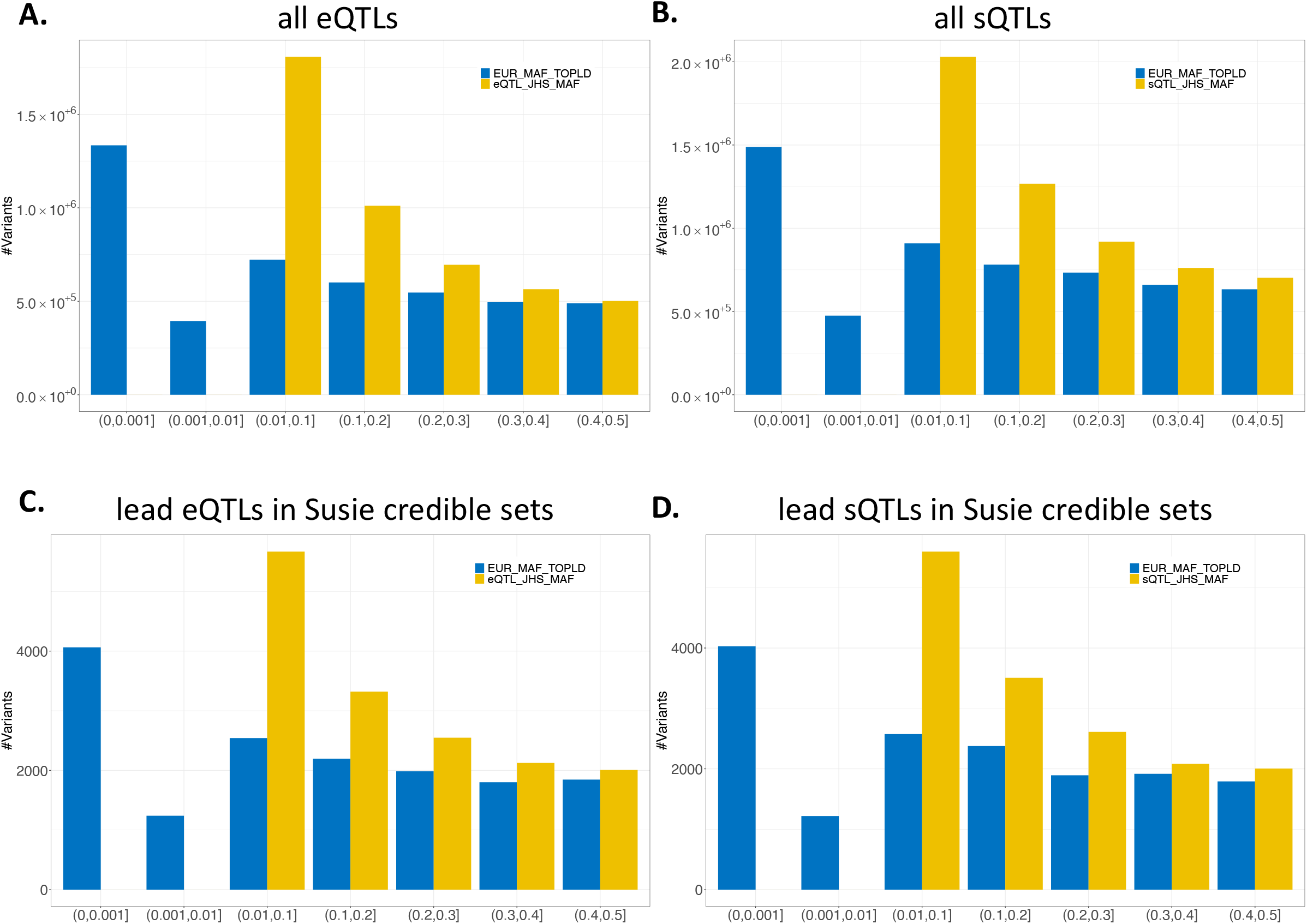
Minor allele frequency of significant QTLs comparison between JHS study samples and European ancestry individuals based on TOP-LD [35504290]. **A and B.** Comparison for all the QTLs. **C and D.** Comparison for lead variants within each SuSie credible set of QTLs. The x-axis is showing MAF bins and the y-axis is showing the counts of variants.

To benefit the scientific community and for easy access to our results, we developed a website for the JHS eQTL/sQTL database (**Figure S6**), where users can search, sort and download our eQTL/sQTL summary statistics without restrictions, as noted below under data availability. In addition, we provide URLs for each variant and gene linking to external sources of variant and gene annotation (e.g., Bravo variant browser, GeneCards, etc.).

In summary, we generated RNA-seq data from PBMC in >1,000 predominantly African ancestry individuals from the JHS study. Leveraging both RNA-seq based gene expression dataset in AFR individuals and WGS data from TOPMed, our QTL analyses reveal many eQTLs and sQTLs missed by previous studies, which either primarily focused on EUR individuals or analyzed AFR data with smaller sample sizes. We also performed fine-mapping for our eQTL and sQTL results and provided credible sets of distinct eQTL and sQTL signals to aid in future colocalization analysis. Our web database provides an easily accessible, convenient interface platform for researchers to search blood-based QTLs among AFR individuals. The large sample size of AFR participants from JHS substantially increases eQTL/sQTL identification power and leads to novel putative mechanisms for GWAS identified variants, demonstrating the need for larger omic data sample sizes across diverse populations, especially as more GWAS are conducted in individuals with significant non-European ancestry. Our results have broad potential to prioritize or validate variants which are more common among African ancestry individuals.

## Data and code availability

Data generated for this study can be accessed via the JHS-QTL web portal: http://jhsqtl.genetics.unc.edu/

## Acknowledgement

The project is supported by funding from the National Institutes of Health, through R01 HL129132 (YL), R01AG075884 (LMR), and R01HL146500 (APR). YL is also partially supported by U01HG011720 and R01HL163972. J.W. was supported by NIH grant 5T32ES007018. The content is solely the responsibility of the authors and does not necessarily represent the official views of the NIH.

We gratefully acknowledge University of Washington Northwest Genomics Center that generated JHS RNA-sequencing data and assisted with appropriate data cleaning and preparation queries. We would also like to thank Stephen Montgomery and Page Goddard for their assistance with the AFGR pre-publication summary statistics.

The JHS is supported and conducted in collaboration with Jackson State University (HHSN268201800013I), Tougaloo College (HHSN268201800014I), the Mississippi State Department of Health (HHSN268201800015I) and the University of Mississippi Medical Center (HHSN268201800010I, HHSN268201800011I and HHSN268201800012I) contracts from the National Heart, Lung, and Blood Institute (NHLBI) and the National Institute on Minority Health and Health Disparities (NIMHD). The authors also wish to thank the staff and participants of the JHS.

## Web resources

JHS-QTL: http://jhsqtl.genetics.unc.edu/

APEX: https://corbinq.github.io/apex/doc/mode_cis/

GCTA: https://yanglab.westlake.edu.cn/software/gcta/#Overview

TOP-LD: http://topld.genetics.unc.edu/

TOPMed: https://topmed.nhlbi.nih.gov/

STAR: https://github.com/alexdobin/STAR

LeafCutter: https://davidaknowles.github.io/leafcutter/

GTEx portal: https://gtexportal.org/home/

Human Protein Atlas portal: proteinatlas.org

AFGR: https://github.com/smontgomlab/AFGR

## References

1. Lappalainen, T., Sammeth, M., Friedländer, M.R., ‘t Hoen, P.A.C., Monlong, J., Rivas, M.A., Gonzàlez-Porta, M., Kurbatova, N., Griebel, T., Ferreira, P.G., et al. (2013). Transcriptome and genome sequencing uncovers functional variation in humans. Nature 501, 506–511.

2. Morley, M., Molony, C.M., Weber, T.M., Devlin, J.L., Ewens, K.G., Spielman, R.S., and Cheung, V.G. (2004). Genetic analysis of genome-wide variation in human gene expression. Nature 430, 743–747.

3. GTEx Consortium (2020). The GTEx Consortium atlas of genetic regulatory effects across human tissues. Science 369, 1318–1330.

4. Li, Y.I., van de Geijn, B., Raj, A., Knowles, D.A., Petti, A.A., Golan, D., Gilad, Y., and Pritchard, J.K. (2016). RNA splicing is a primary link between genetic variation and disease. Science 352, 600–604.

5. Võsa, U., Claringbould, A., Westra, H.-J., Bonder, M.J., Deelen, P., Zeng, B., Kirsten, H., Saha, A., Kreuzhuber, R., Yazar, S., et al. (2021). Large-scale cis- and trans-eQTL analyses identify thousands of genetic loci and polygenic scores that regulate blood gene expression. Nat. Genet. 53, 1300–1310.

6. Stranger, B.E., Montgomery, S.B., Dimas, A.S., Parts, L., Stegle, O., Ingle, C.E., Sekowska, M., Smith, G.D., Evans, D., Gutierrez-Arcelus, M., et al. (2012). Patterns of cis regulatory variation in diverse human populations. PLoS Genet. 8, e1002639.

7. Shang, L., Smith, J.A., Zhao, W., Kho, M., Turner, S.T., Mosley, T.H., Kardia, S.L.R., and Zhou, X. (2020). Genetic Architecture of Gene Expression in European and African Americans: An eQTL Mapping Study in GENOA. Am. J. Hum. Genet. 106, 496–512.

8. Mogil, L.S., Andaleon, A., Badalamenti, A., Dickinson, S.P., Guo, X., Rotter, J.I., Johnson, W.C., Im, H.K., Liu, Y., and Wheeler, H.E. (2018). Genetic architecture of gene expression traits across diverse populations. PLoS Genet. 14, e1007586.

9. Mak, A.C.Y., Kachuri, L., Hu, D., Eng, C., Huntsman, S., Elhawary, J.R., Gupta, N., Gabriel, S., Xiao, S., Gui, H., et al. (2021). Gene expression in African Americans and Latinos reveals ancestry-specific patterns of genetic architecture. BioRxiv.

10. Wilson, J.G., Rotimi, C.N., Ekunwe, L., Royal, C.D.M., Crump, M.E., Wyatt, S.B., Steffes, M.W., Adeyemo, A., Zhou, J., Taylor, H.A., et al. (2005). Study design for genetic analysis in the Jackson Heart Study. Ethn. Dis. 15, S6–30.

11. Jun, G., Flickinger, M., Hetrick, K.N., Romm, J.M., Doheny, K.F., Abecasis, G.R., Boehnke, M., and Kang, H.M. (2012). Detecting and estimating contamination of human DNA samples in sequencing and array-based genotype data. Am. J. Hum. Genet. 91, 839– 848.

12. Conomos, M.P., Miller, M.B., and Thornton, T.A. (2015). Robust inference of population structure for ancestry prediction and correction of stratification in the presence of relatedness. Genet. Epidemiol. 39, 276–293.

13. Li, Y.I., Knowles, D.A., Humphrey, J., Barbeira, A.N., Dickinson, S.P., Im, H.K., and Pritchard, J.K. (2018). Annotation-free quantification of RNA splicing using LeafCutter. Nat. Genet. 50, 151–158.

14. Quick, C., Guan, L., Li, Z., Li, X., Dey, R., Liu, Y., Scott, L., and Lin, X. (2020). A versatile toolkit for molecular QTL mapping and meta-analysis at scale. BioRxiv.

15. Yang, J., Benyamin, B., McEvoy, B.P., Gordon, S., Henders, A.K., Nyholt, D.R., Madden, P.A., Heath, A.C., Martin, N.G., Montgomery, G.W., et al. (2010). Common SNPs explain a large proportion of the heritability for human height. Nat. Genet. 42, 565–569.

16. Yang, J., Lee, S.H., Goddard, M.E., and Visscher, P.M. (2011). GCTA: a tool for genome-wide complex trait analysis. Am. J. Hum. Genet. 88, 76–82.

17. Stegle, O., Parts, L., Piipari, M., Winn, J., and Durbin, R. (2012). Using probabilistic estimation of expression residuals (PEER) to obtain increased power and interpretability of gene expression analyses. Nat. Protoc. 7, 500–507.

18. Wang, G., Sarkar, A., Carbonetto, P., and Stephens, M. (2020). A simple new approach to variable selection in regression, with application to genetic fine mapping. J. Royal Statistical Soc. B.

19. Rowland, B., Huh, R., Hou, Z., Crowley, C., Wen, J., Shen, Y., Hu, M., Giusti-Rodríguez, P., Sullivan, P.F., and Li, Y. (2022). THUNDER: A reference-free deconvolution method to infer cell type proportions from bulk Hi-C data. PLoS Genet. 18, e1010102.

20. Barbeira, A.N., Bonazzola, R., Gamazon, E.R., Liang, Y., Park, Y., Kim-Hellmuth, S., Wang, G., Jiang, Z., Zhou, D., Hormozdiari, F., et al. (2021). Exploiting the GTEx resources to decipher the mechanisms at GWAS loci. Genome Biol. 22, 49.

21. Sun, Q., Crowley, C.A., Huang, L., Wen, J., Chen, J., Bao, E.L., Auer, P.L., Lettre, G., Reiner, A.P., Sankaran, V.G., et al. (2022). From GWAS variant to function: A study of □ 148,000 variants for blood cell traits. HGG Adv. 3, 100063.

22. Chen, M.-H., Raffield, L.M., Mousas, A., Sakaue, S., Huffman, J.E., Moscati, A., Trivedi, B., Jiang, T., Akbari, P., Vuckovic, D., et al. (2020). Trans-ethnic and Ancestry-Specific Blood-Cell Genetics in 746,667 Individuals from 5 Global Populations. Cell 182, 1198–1213.e14.

23. Huang, L., Rosen, J.D., Sun, Q., Chen, J., Wheeler, M.M., Zhou, Y., Min, Y.-I., Kooperberg, C., Conomos, M.P., Stilp, A.M., et al. (2022). TOP-LD: A tool to explore linkage disequilibrium with TOPMed whole-genome sequence data. Am. J. Hum. Genet. 109, 1175–1181.

24. Gautam, Y., Caldwell, J., Kottyan, L., Chehade, M., Dellon, E.S., Rothenberg, M.E., Mersha, T.B., and Consortium of Eosinophilic Gastrointestinal Disease Researchers (CEGIR) investigators (2022). Genome-wide admixture and association analysis identifies African ancestry-specific risk loci of eosinophilic esophagitis in African Americans. J. Allergy Clin. Immunol.

