## Supplemental Materials for "Gene Expression and Splicing QTL Analysis of Blood Cells in African American Participants from the Jackson Heart Study"

### The Jackson Heart Study

The Jackson Heart Study (JHS) is a community-based cohort study investigating causes of cardiovascular and related diseases among African Americans with the aim of more effective prevention and treatment. JHS recruited 5,306 participants from the Jackson, Mississippi metropolitan area [1]; PBMCs from a subset of consenting participants were cryopreserved, as described in Wilson et al [1]. Whole-genome sequencing data from 3,406 JHS participants are available from the NHLBI Trans-Omics for Precision Medicine (TOPMed) Program [2]; we here use whole genome sequencing data with minimum depth 0 (DP0) from freeze 8 (unphased), with methods described at https://topmed.nhlbi.nih.gov/topmed-whole-genome-sequencing-methods-freeze-8.

### Ancestry background of JHS samples

We performed local and global ancestry inference using RFMix [3] for the 1,020 JHS participants used in the analyses, with reference panels from a combination of HGDP for Native American populations and 1000G for Europeans, Africans, East Asians and South Asians. Specifically, we constructed a balanced reference panel with each consisting of 92 individuals to match with the smallest sample size. For 1000G, we classified CEU, TSI, FIN, GBR, and IBS as Europeans; YRI, LWK, GWD, MSL, ESN, ASW and ACB as Africans; CHB, JPT, CHS, CDX, KHV as East Asians; and GIH, PJL, BEB, STU, ITU as South Asians. The mean and median African global ancestry component is 83.4% and 85.4% (minimum 26.0% and maximum 98.1%).

### RNA-seq quantification

RNA sequencing for all samples is performed at the University of Washington Northwest Genomics Center (NWGC) with an average read-depth of 50M. For library construction, total RNA was scaled to 7.5ng/ul (total volume of 50ul) on the Perkin Elmer Janus II Workstation. Poly-A selection and cDNA synthesis were performed using the TruSeq Stranded mRNA kit (Illumina, cat#RS-122-2103). All steps were automated on the Perkin Elmer Sciclone NGSx Workstation. Final RNASeq libraries were quantified using the Quant-it dsDNA High Sensitivity assay, and library insert size distribution was checked using a fragment analyzer (Advanced Analytical; kit ID DNF474). Samples where adapter dimers constituted more than 4% of the electopherogram area were excluded. Successful libraries were normalized and pooled prior to sequencing. For read processing, base call generation was conducted on the NovaSeq6000 instrument (RTA 3.1.5). Next, demultiplexed, unaligned BAM files produced by Picard ExtractIlluminaBarcodes and IlluminaBasecallsToSam were converted to FASTQ format using SamTools bam2fq (v1.4). Sequence read and base quality checking was performed using the FASTX-toolkit (v0.0.13), and sequence alignment to GRCh38 with reference transcriptome GENCODE release 30 was performed using STAR (v2.6.1d) [4].

**Quality control**

RNA sequencing data were generated for a total of 1,027 JHS African ancestry individuals in the University of Washington Northwest Genomics Center (NWGC) with an average read-depth of 50M. The gene-level expression quantification was estimated as transcripts per million (TPM) for each gene, which was quantitated and released on the merged-level data across individuals with RSEM (v1.3.1). We then performed sample-, variant- and gene-level QC for the RNA-seq data. We removed 5 individuals due to low RNA-seq quality and 2 individuals due to the mismatched sex compared to the recorded sex from JHS. We then used VerifyBamID to check the individual consistency between RNA-seq and WGS data [5]. 8 individuals who failed VerifyBamID (inconsistency rate >= 2%), likely due to potential mixing or contamination of the samples, were also removed. After sample-level QC, a total of 1,012 predominantly African ancestry individuals were left for eQTL and sQTL analysis. We further performed variant- and gene-level QC after removing these individuals. Variants with minor allele frequency (MAF) > 1% in our 1,012 QC+ samples were kept in further analysis. For gene-level QC, we followed a similar strategy as applied in the GTEx project [6]: genes were kept with expression ≥ 0.1 transcripts per million (TPM), expressed in at least 20% of individuals, and having ≥ 6 expected read counts from RSEM in at least 20% of individuals. After gene-level QC, we in total have 17,383 genes left for eQTL analysis. We used PC-AiR [7] to obtain genetic PCs accounting for genetic relatedness in JHS, and used the top 10 genotype PCs in QTL analyses. We applied an aggressive search of potential confounders with controlling 10-130 probabilistic estimations of expression residuals (PEER) factors [8], leading to selection of 70 PEER factors, where we no longer observed a substantive increase in number of detected QTLs with the addition of an additional 10 PEER factors (**Figure S4A**).

In the sQTL analysis, we first converted aligned BAM files to Fastq files using Picard v2.26.11 and re-aligned the reads to GRCh38 reference genome using STAR (v2.7.0a) [4]. For the re-alignment, we obtained individual-level genotypes from TOPMed freeze 8 WGS data and used that in the WASP option of STAR to avoid allelic mapping bias that would affect sQTL results [4,9]; this is why mapping had to redone versus the standard STAR mapping described above in “RNA-seq quantification**”**. After realignment, we calculated intron excision ratios to quantify alternative splicing events using LeafCutter [10]. We first identified intron clusters by excision ratios and followed a similar QC strategy as implemented in the GTEx project [6], including at least 30 split reads supporting each intron cluster, at least 0.001 of cluster ratio for each intron cluster and up to 500kb of introns at this step. After initial QC, we mapped the intron clusters to genes with GENCODE release version 38. Furthermore, introns with no read counts in more than 50% of samples, with low complexity and low variability across samples were also removed [6,10]. Finally, 97,840 intron clusters mapped to genes were left in our sQTL analysis. Similar to PEER factor selection for our eQTL analysis, we searched for the optimal number of splicing PCs in the range of 10-50, leading to selection of 25 splicing PCs, which led to the largest number of significant variant-cluster pairs (**Figure S4B**).

**Fine mapping**
We performed fine mapping from eQTL or sQTL summary statistics using SuSie [11]. We calculated the in-sample LD using plink [12] from the 1,012 JHS samples used in QTL discoveries. We first set the maximum allowed independent signals to be 10, and then increased the threshold one at a time if the number of credible sets hit the threshold. We only performed fine-mapping analyses for genes (or gene-clusters) with top eQTL (or sQTL) p-value < 5e-8 to ensure model stability.

**Web design**
We developed a website for the JHS eQTL/sQTL results (http://jhsqtl.genetics.unc.edu) (**Figure S6**). This website was built using the LAMP (Linux, Apache2, MySQL, and PHP) framework. For dataset preparation, we selected eQTL-gene pairs or sQTL-gene cluster pairs at FDR 5% and added rsID for each variant using NCBI dbSNP Build 155. For visualization, we applied datatable.js to present the eQTLs (4,798,604 total rows) and sQTLs (5,921,368 total rows) separately. We enabled users to search, sort, and download QTL results without restriction. In addition, we provided URLs to external sources (e.g., link to bravo for variants and link to Genecard for genes) to provide additional information.

**Supplemental acknowledgement**
The Genotype-Tissue Expression (GTEx) Project was supported by the Common Fund of the Office of the Director of the National Institutes of Health, and by NCI, NHGRI, NHLBI, NIDA, NIMH, and NINDS. The data (release 8) used for the analyses described in this manuscript were obtained from the GTEx Portal on 06/20/2022.

Molecular data for the Trans-Omics in Precision Medicine (TOPMed) program was supported by the National Heart, Lung and Blood Institute (NHLBI). Genome sequencing for “NHLBI TOPMed: The Jackson Heart Study” (phs000964.v1.p1) was performed at the Northwest Genomics Center (HHSN268201100037C). Core support including centralized genomic read mapping and genotype calling, along with variant quality metrics and filtering were provided by the TOPMed Informatics Research Center (3R01HL-117626-02S1; contract HHSN268201800002I). Core support including phenotype harmonization, data management, sample-identity QC, and general program coordination were provided by the TOPMed Data Coordinating Center (R01HL-120393; U01HL-120393; contract HHSN268201800001I). We gratefully acknowledge the studies and participants who provided biological samples and data for TOPMed. The TOPMed Banner Authorship List can be found at: https://www.nhlbiwgs.org/topmed-banner-authorship.

### Supplemental Figures

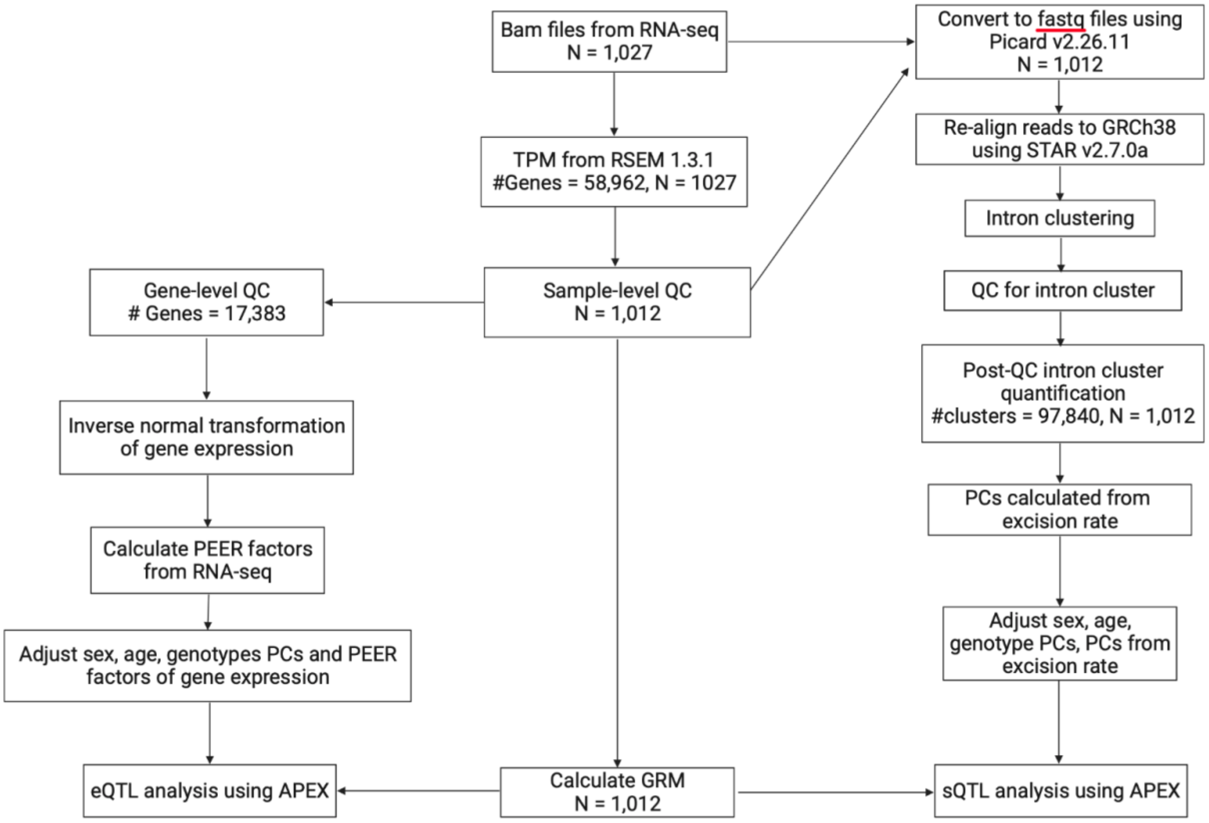


**Figure S1. JHS eQTL and sQTL analysis workflow.** PCs: principal components; GRM: genetic relationship matrix; QC: quality control; sQTL: splicing quantitative trait loci; APEX: All-in-one Package for Efficient Xqtl analysis.

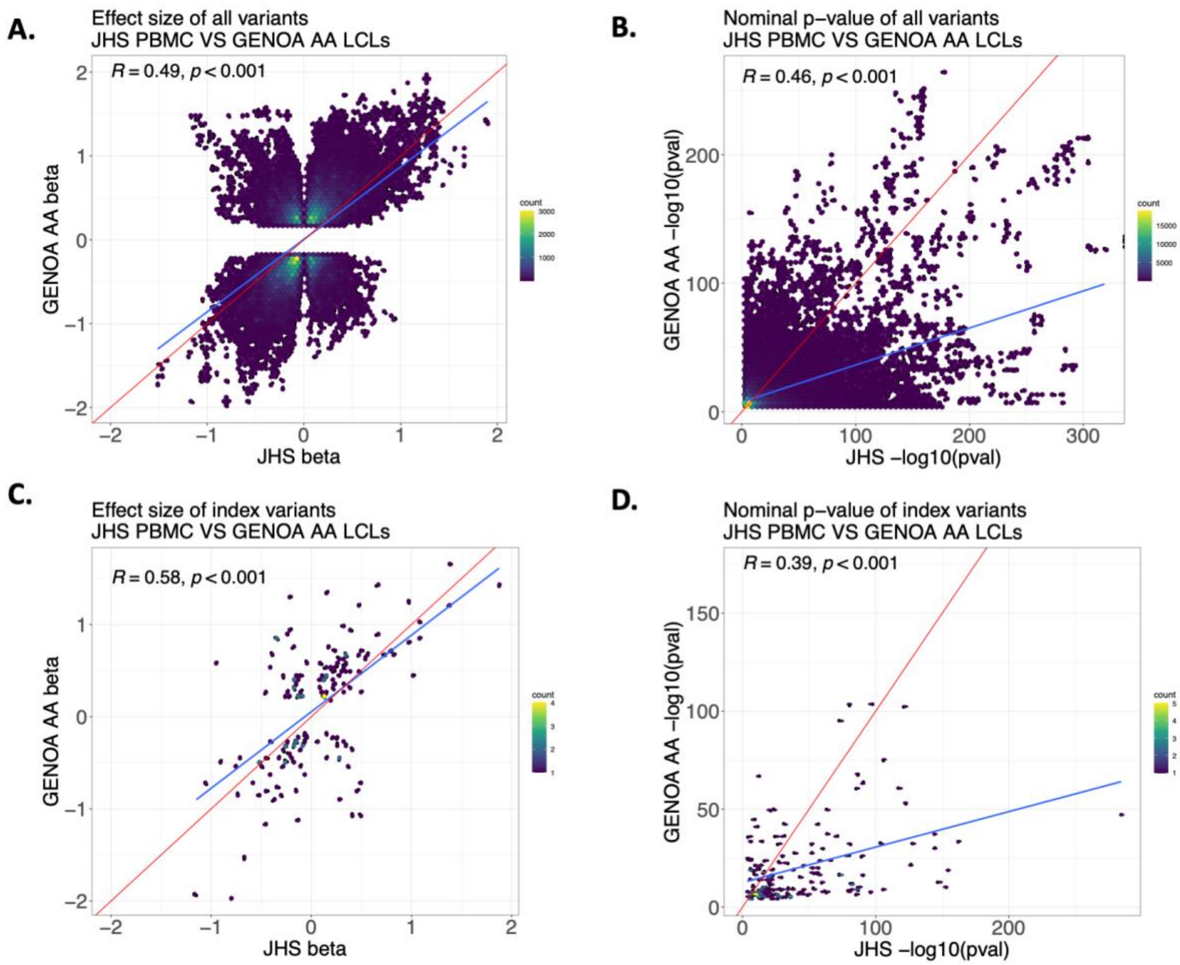


**Figure S2. eQTL results comparison with GENOA LCL eQTLs. A and B.** Comparison of effect size estimates (**A**) and nominal p-values (**B**), for all shared variants. **C and D.** Comparison of effect size estimates (**C**) and nominal p-values (**D**) restricting to the index variants (i.e., the most significant eQTL variant for each gene). The red line denotes the diagonal line and the blue line is the regression fitting line.

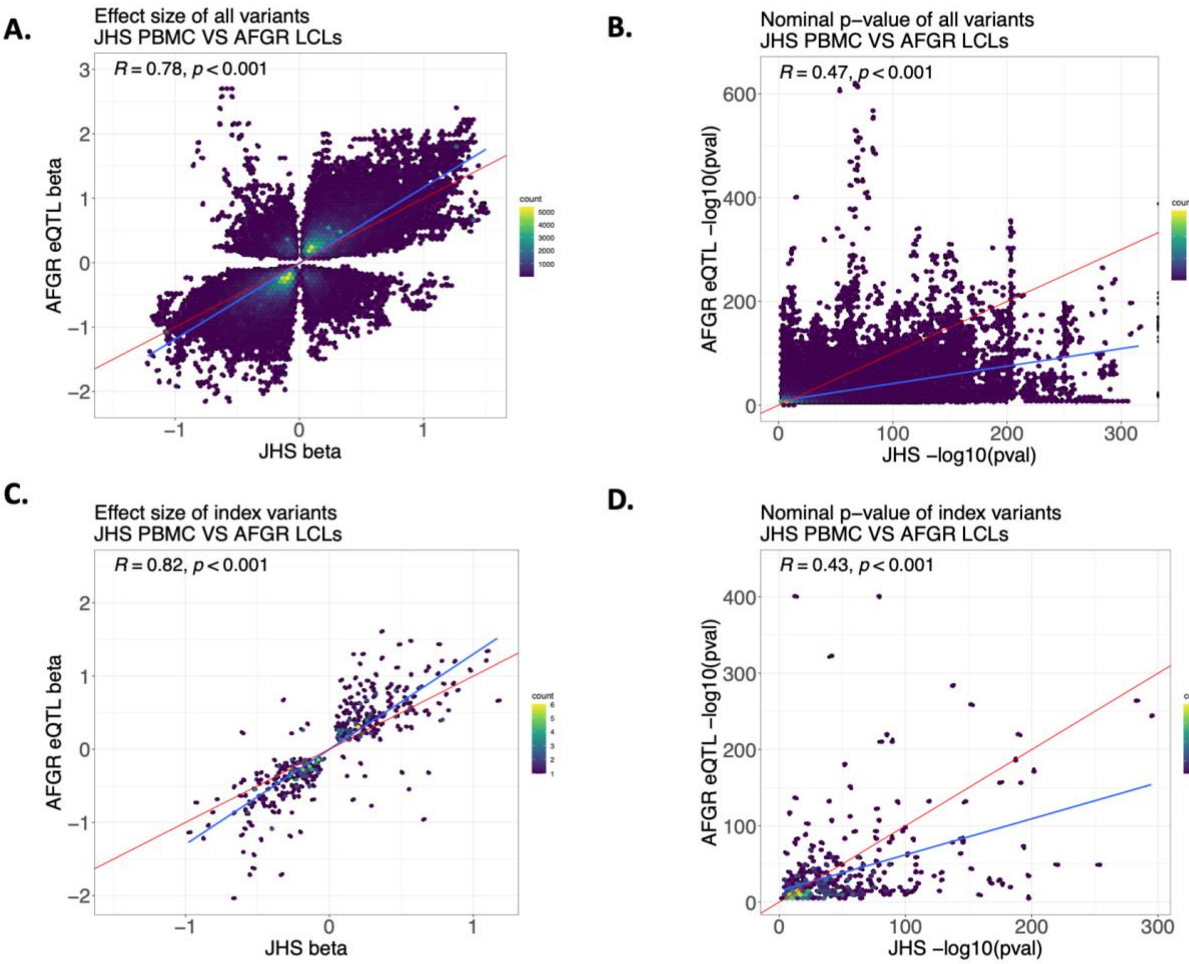


**Figure S3. eQTL results comparison with AFGR pre-publication eQTL. A and B.** Comparison of effect size estimates (**A**) and nominal p-values (**B**), for all shared variants. **C and D.** Comparison of effect size estimates (**C**) and nominal p-values (**D**) restricting to the index variants (i.e., the most significant eQTL variant for each gene). The red line denotes the diagonal line and the blue line is the regression fitting line.

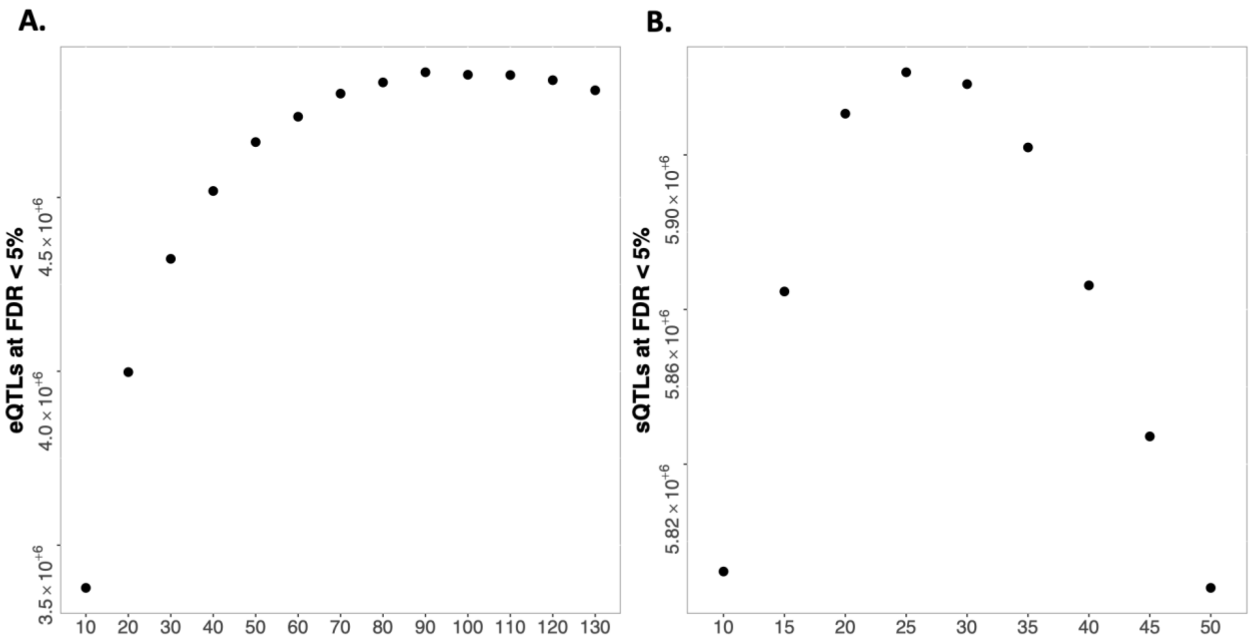


**Figure S4. Choice of the numbers of splicing PCs and PEER factors. A**. Number of PEER factors (X axis) versus the number of significant (FDR < 5%) eQTLs (Y axis). **B**. Number of Splicing PCs (X axis) versus the number of significant (FDR < 5%) sQTLs (Y axis).

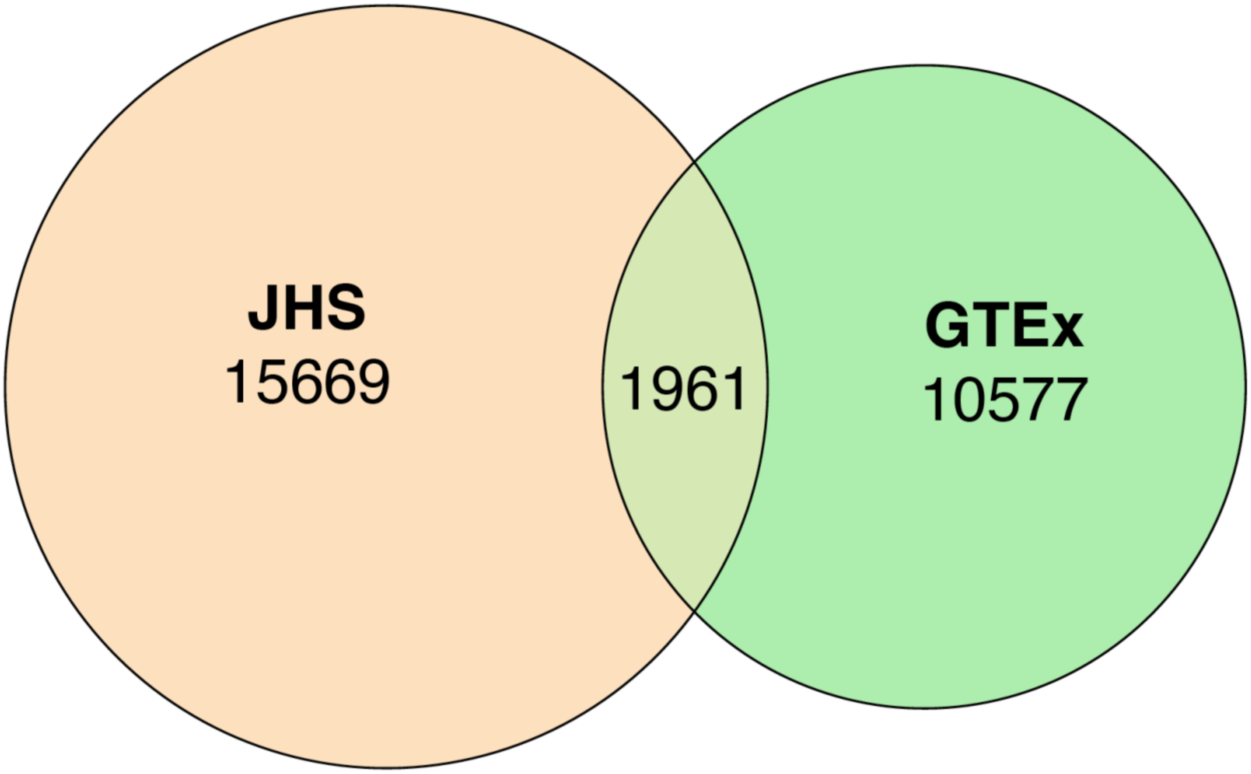


**Figure S5. Venn diagram to show the independent eQTL signals comparison between the JHS credible sets GTEx distinct signals** [32913098]**.** The comparison was restricted to 6,779 genes where we performed fine-mapping and that also have at least one distinct signal in GTEx. Signals are listed as overlapping if the GTEx distinct signal is in JHS credible sets.

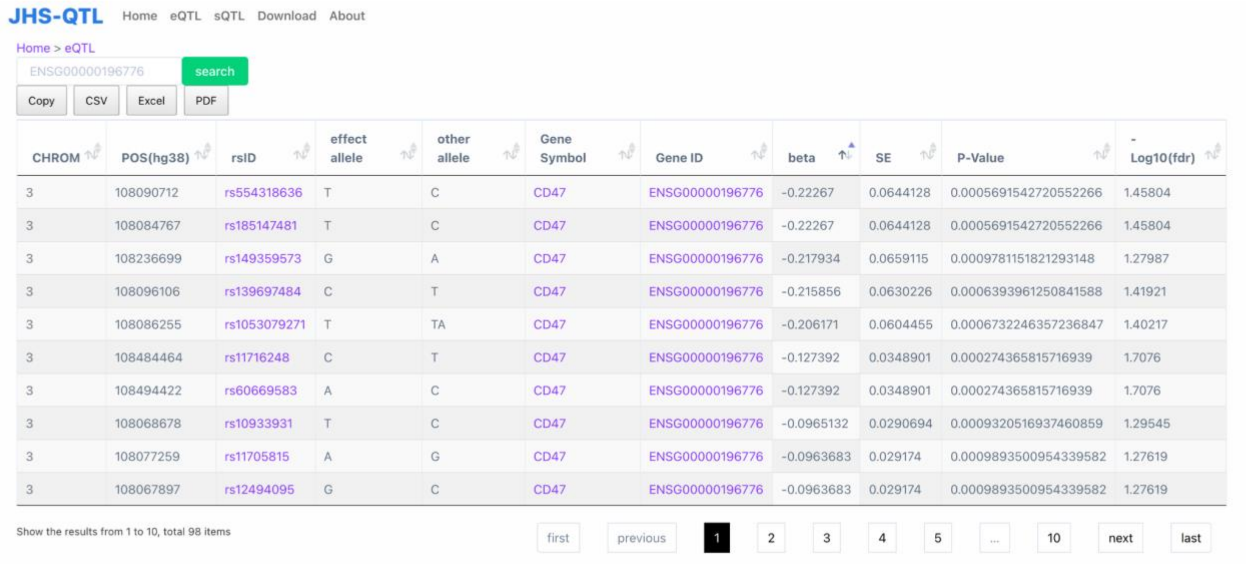


| CHROM | POS(hg38) | rsID | effect allele | other allele | Gene Symbol | Gene ID | beta | SE | P-Value | -Log10(fdr) |
|---|---|---|---|---|---|---|---|---|---|---|
| 3 | 108090712 | rs554318636 | T | C | CD47 | ENSG00000196776 | -0.22267 | 0.0644128 | 0.0005691542720552266 | 1.45804 |
| 3 | 108084767 | rs185147481 | T | C | CD47 | ENSG00000196776 | -0.22267 | 0.0644128 | 0.0005691542720552266 | 1.45804 |
| 3 | 108236699 | rs149359573 | G | A | CD47 | ENSG00000196776 | -0.217934 | 0.0659115 | 0.0009781151821293148 | 1.27987 |
| 3 | 108096106 | rs139697484 | C | T | CD47 | ENSG00000196776 | -0.215856 | 0.0630226 | 0.0006393961250841588 | 1.41921 |
| 3 | 108086255 | rs1053079271 | T | TA | CD47 | ENSG00000196776 | -0.206171 | 0.0604455 | 0.0006732246357236847 | 1.40217 |
| 3 | 108484464 | rs11716248 | C | T | CD47 | ENSG00000196776 | -0.127392 | 0.0348901 | 0.000274365815716939 | 1.7076 |
| 3 | 108494422 | rs60669583 | A | C | CD47 | ENSG00000196776 | -0.127392 | 0.0348901 | 0.000274365815716939 | 1.7076 |
| 3 | 108068678 | rs10933931 | T | C | CD47 | ENSG00000196776 | -0.0965132 | 0.0290694 | 0.0009320516937460859 | 1.29545 |
| 3 | 108077259 | rs11705815 | A | G | CD47 | ENSG00000196776 | -0.0963683 | 0.029174 | 0.0009893500954339582 | 1.27619 |
| 3 | 108067897 | rs12494095 | G | C | CD47 | ENSG00000196776 | -0.0963683 | 0.029174 | 0.0009893500954339582 | 1.27619 |



**Figure S6. An example of JHS eQTL query results.** The example shows eQTLs results for gene *CD47*.

### References


1. Wilson, J.G., Rotimi, C.N., Ekunwe, L., Royal, C.D.M., Crump, M.E., Wyatt, S.B., Steffes, M.W., Adeyemo, A., Zhou, J., Taylor, H.A., et al. (2005). Study design for genetic analysis in the Jackson Heart Study. Ethn. Dis. *15*, S6-30.

2. Taliun, D., Harris, D.N., Kessler, M.D., Carlson, J., Szpiech, Z.A., Torres, R., Taliun, S.A.G., Corvelo, A., Gogarten, S.M., Kang, H.M., et al. (2021). Sequencing of 53,831 diverse genomes from the NHLBI TOPMed Program. Nature *590*, 290–299.

3. Maples, B.K., Gravel, S., Kenny, E.E., and Bustamante, C.D. (2013). RFMix: a discriminative modeling approach for rapid and robust local-ancestry inference. Am. J. Hum. Genet. *93*, 278–288.

4. Dobin, A., Davis, C.A., Schlesinger, F., Drenkow, J., Zaleski, C., Jha, S., Batut, P., Chaisson, M., and Gingeras, T.R. (2013). STAR: ultrafast universal RNA-seq aligner. Bioinformatics *29*, 15–21.

5. Jun, G., Flickinger, M., Hetrick, K.N., Romm, J.M., Doheny, K.F., Abecasis, G.R., Boehnke, M., and Kang, H.M. (2012). Detecting and estimating contamination of human DNA samples in sequencing and array-based genotype data. Am. J. Hum. Genet. *91*, 839–848.

6. GTEx Consortium (2020). The GTEx Consortium atlas of genetic regulatory effects across human tissues. Science *369*, 1318–1330.

7. Conomos, M.P., Miller, M.B., and Thornton, T.A. (2015). Robust inference of population structure for ancestry prediction and correction of stratification in the presence of relatedness. Genet. Epidemiol. *39*, 276–293.

8. Stegle, O., Parts, L., Piipari, M., Winn, J., and Durbin, R. (2012). Using probabilistic estimation of expression residuals (PEER) to obtain increased power and interpretability of gene expression analyses. Nat. Protoc. *7*, 500–507.

9. van de Geijn, B., McVicker, G., Gilad, Y., and Pritchard, J.K. (2015). WASP: allele-specific software for robust molecular quantitative trait locus discovery. Nat. Methods *12*, 1061–1063.

10. Li, Y.I., Knowles, D.A., Humphrey, J., Barbeira, A.N., Dickinson, S.P., Im, H.K., and Pritchard, J.K. (2018). Annotation-free quantification of RNA splicing using LeafCutter. Nat. Genet. *50*, 151–158.

11. Weissbrod, O., Hormozdiari, F., Benner, C., Cui, R., Ulirsch, J., Gazal, S., Schoech, A.P., van de Geijn, B., Reshef, Y., Márquez-Luna, C., et al. (2020). Functionally informed fine-mapping and polygenic localization of complex trait heritability. Nat. Genet. *52*, 1355–1363.

12. Purcell, S., Neale, B., Todd-Brown, K., Thomas, L., Ferreira, M.A.R., Bender, D., Maller, J., Sklar, P., de Bakker, P.I.W., Daly, M.J., et al. (2007). PLINK: a tool set for whole-genome association and population-based linkage analyses. Am. J. Hum. Genet. *81*, 559–575.
